# Assessment of the Viscoelastic Mechanical Properties of the Porcine Optic Nerve Head using Micromechanical Testing and Finite Element Modeling

**DOI:** 10.1101/2021.04.16.440170

**Authors:** Babak N. Safa, A. Thomas Read, C. Ross Ethier

**Affiliations:** Wallace H. Coulter Department of Biomedical Engineering, Georgia Institute of Technology/Emory University, Atlanta GA, USA

**Author notes:** **Corresponding author:** Petit Biotechnology Building (IBB), 315 Ferst Drive, Room 2306, Atlanta, GA 30332-0363.

**Keywords:** Optic Nerve Head, Glaucoma, Porcine Eye, Incremental Creep, Mechanical Stress, Viscoelasticity

## Abstract

Optic nerve head (ONH) biomechanics is centrally involved in the pathogenesis of glaucoma, a blinding ocular condition often characterized by elevation and fluctuation of the intraocular pressure and resulting loads on the ONH. Further, tissue viscoelasticity is expected to strongly influence the mechanical response of the ONH to mechanical loading, yet the viscoelastic mechanical properties of the ONH remain unknown. To determine these properties, we conducted micromechanical testing on porcine ONH tissue samples, coupled with finite element modeling based on a mixture model consisting of a biphasic material with a viscoelastic solid matrix. Our results provide a detailed description of the viscoelastic properties of the porcine ONH at each of its four anatomical quadrants (i.e., nasal, superior, temporal, and inferior). We showed that the ONH’s viscoelastic mechanical response can be explained by a dual mechanism of fluid flow and solid matrix viscoelasticity, as is common in other soft tissues. We obtained porcine ONH properties as follows: matrix Young’s modulus *E*=1.895 [1.056,2 .391] kPa (median [min., max.]), Poisson’s ratio *ν*=0.142 [0.060,0 .312], kinetic time-constant *τ*=214 [89,921] sec, and hydraulic permeability *k*=3.854 × 10^−1^ [3.457 × 10^−2^,9.994 × 10^−1^] mm^4^/(N sec). These values can be used to design and fabricate physiologically appropriate *ex vivo* test environments (e.g., 3D cell culture) to further understand glaucoma pathophysiology.

## 1 Introduction

Glaucoma is the leading cause of irreversible blindness, with c. 78 million patients worldwide (National Eye Institute, 2012; World Health Organization, 2019). Glaucoma is an optic neuropathy characterized by chronic deterioration of vision due to gradual loss of retinal ganglion cells (RGC) (Weinreb et al., 2014). The most common form of glaucoma is primary open-angle glaucoma (POAG), which is strongly associated with elevation of the intraocular pressure (IOP), thought to cause biomechanical insult to the RGC axons and the glial cells of the optic nerve head (ONH). This results in remodeling and often irreversible posterior deformation of the ONH (cupping) (Weinreb et al., 2014). Therefore, ONH biomechanics is a key aspect of POAG pathophysiology. It is known that IOP fluctuates significantly (Abelson, 2011; Kida et al., 2006; Turner et al., 2019), yet little is known about the viscoelastic mechanical behavior of the ONH.

Many soft tissues behave differently under static vs. dynamic loading (i.e., viscoelasticity), due to their high water content and the intrinsic viscoelastic behavior of the extracellular matrix (ECM) (Connizzo and Grodzinsky, 2017; Oftadeh et al., 2018). Tissue viscoelasticity strongly influences the magnitude and frequency of the mechanical loads on cells resident in the tissue, i.e. it influences the mechanobiological signals the cells receive from their environment. Further, fluid-dependent viscoelasticity (also known as poroelastic viscoelasticity) plays a key role in material transport (e.g., cellular nutrients and waste) to and from the cells. Therefore, viscoelasticity is an important aspect of understanding cell mechanobiology, which influences tissue remodeling (Martino et al., 2018).

We previously showed significant viscoelastic effects in the porcine and murine ONHs, as evident from a significant dependence of the biomechanical response on mechanical loading rate (Boazak et al., 2019). Additionally, Myers and co-workers showed that the posterior bovine sclera, a region of the ocular globe that includes the ONH, shows a nonlinear and viscoelastic mechanical response (Myers et al., 2010). However, no studies report the viscoelastic material properties of the ONH.

We hypothesized that the ONH’s mechanical response could be explained using mixture theory based on a biphasic model with an inherently viscoelastic matrix. Therefore, this study’s objective was to assess the viscoelastic mechanical properties of the porcine ONH using micromechanical testing and finite element modeling. We chose to study the porcine ONH because porcine eyes are readilyavailable and have been previously used to study glaucoma (Ruiz-Ederra et al., 2005); because porcine ocular anatomy is similar to human anatomy, particularly in the ONH possessing a collagenous lamina cribrosa (Gogola et al., 2018; Sanchez et al., 2011); and because the porcine optic disc size is similar to that in the human, making it appropriate for mechanical testing (Boazak et al., 2019). Our results provide a detailed characterization of the mechanical properties of the ONH, which in turn provides new insight into the mechanophysiology of the ONH and further elucidates the role of ONH biomechanics in glaucoma.

## 2 Methods

### 2.1 Experimental sample preparation

For mechanical analyses of the ONH, 8 eyes were collected from 4 freshly sacrificed female pigs (Domestic Yorkshire Crossbred Swine [common farm pig in the US], c. 5 months of age) by resection from the orbital cavity either immediately or within a few hours after death. Careful incisions were made through the extraocular muscles, the surrounding fascia, and the optic nerve to isolate the globe while keeping portions of the surrounding lid and skin attached, which were used as anatomical references (Figure 1A).

**Figure 1:**
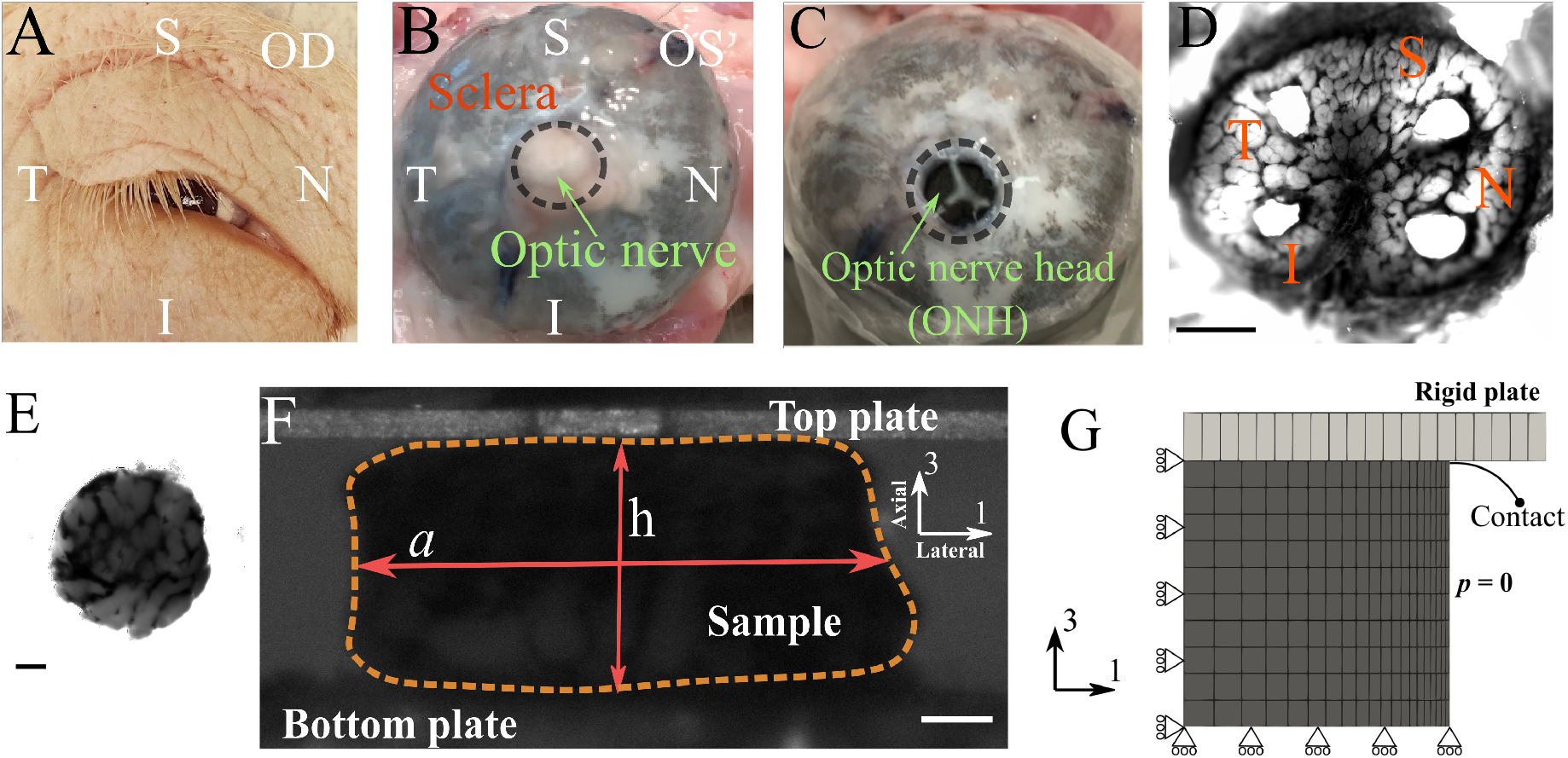
Overview of the experimental procedure used in this study. (A) an example of the harvested eye [OD] and surrounding tissue with labeling for anatomical orientation [I=inferior, N=nasal, S=superior, T=temporal]. (B) posterior segment of an eyeball [OS] showing the four anatomical quadrants. (C) an eye with the optic nerve transected flush with the posterior scleral surface. (D) A tangential slice of an ONH (scale bar = 1 mm), from which four samples (E; scale bar = 0.1mm) are harvested (one from each quadrant). (F) Each sample was tested using a micromechanical compression testing system, and the height [h] and width [a] of the sample were measured throughout the test (scale bar = 0.1 mm). From these measurements, the axial and lateral strains (engineering strains) were calculated as *ϵ*_axial_ =h/h_0_-1 and *ϵ*_lateral_ =a/a_0_-1, respectively, and the apparent Poisson’s ration was computed as ν_app_ = - *ϵ*_lateral_/*ϵ*_axial_,. (G) An axisymmetric finite element model was used to model the mechanical response of the ONH, where a rigid plate compresses the ONH sample in the axial (axis-3) direction. Rolling boundary conditions were utilized at the bottom and center-line to enforce model symmetry. The lateral face of the sample was prescribed to have zero pressure, which allowed outflow of fluid during compression.

The optic nerve head (ONH) was then dissected (Boazak et al., 2019). Briefly, the ONH was exposed by transection of the optic nerve flush with the sclera’s posterior aspect using a surgical blade (Figure 1 B&C). A 1 mm thick tangential slice posterior to this surface, and encompassing the peripapillary sclera, was then harvested by means of a scalpel blade (Figure 1D). From each anatomical quadrant (i.e., inferior [I], nasal [N], superior [S], and temporal [T]), one cylindrical sample was collected using a 1 mm diameter surgical biopsy punch and placed in ice-cold physiological phosphate buffered saline solution (PBS). Each ONH sample’s cross-sectional image was acquired using bright-field microscopy (Leica DM6; Wetzlar, Germany), which was used later to measure sample diameter (Figure 1E). The samples were then immediately studied via micromechanical testing.

### 2.2 Micromechanical incremental creep test

To measure the viscoelastic mechanical properties of the ONH, we conducted a dynamic mechanical test on each sample using a micromechanical testing device (CellScale Microsquisher, Waterloo, ON, Canada). This instrument measures the deflection of a tungsten cantilever beam with known diameter (0.5508 mm), length (54-57 mm), and elastic properties (Young’s modulus 411 GPa) as the tissue sample is compressed by the beam (Boazak et al., 2019; Brown et al., 2021). During mechanical testing, samples were bathed in PBS at 37°C.

Samples were subjected to a force-controlled incremental creep protocol under unconfined compression between two rigid plates (Figure 1F). The sample was gradually loaded to 200 μN (tare-load) over 60 seconds, immediately unloaded to zero force at the same rate, and then loaded to tare-load once more, after which the tare load was maintained for one minute (recovery phase) to establish a consistent reference testing condition. The sample was imaged throughout the test using a CCD camera (Figure 1F). The test protocol consisted of four incremental loading cycles to 524, 1047, 1470, and 2094 μN. At each force level, the sample was held for 5 min to allow creep deformation. Following the fourth creep cycle, the sample was unloaded to the tare-load level over 4 minutes and held there for 1 minute, which concluded the micromechanical test.

### 2.3 Deformation analyses

We used the height of the sample during the test to calculate axial engineering strain as *ϵ_axial_*=*h/h*_0_−1, where *h* was the height of the sample during the test, and *h_0_* was the reference height of the sample, i.e. the height with the tare load on the sample. The sample’s reference height was measured by averaging the final 20 seconds of the recovery phase before beginning the cycles of creep loading (*h_0_* = 0.5 ± 0.2 mm). To measure the lateral strain, we used image analysis. Specifically, we created a binarized mask that covered the sample, calculated the mask’s area, and calculated the sample’s average lateral width by dividing this area by the height of the mask (Figure 1F). The lateral engineering strain was then calculated as *ϵ_lateral_*=*a/a*_0_−1, where *a* was the average lateral width of the sample and *a_0_* was its reference value, calculated as for *h_0_*. Throughout the test, we also calculated the apparent Poisson’s ratio ν_app_ = **- *ϵ***_lateral_/***ϵ***_axial_, which was used as measure of volumetric compressibility, with ν_app_ = 0.5 indicating incompressibility under the assumption of cylindrical axisymmetry.

### 2.4 Anatomical region heterogeneity in deformation responses

We compared the samples’ mechanical response from the four anatomical regions to determine whether the ONH’s heterogeneous structure affected the outcomes. Specifically, we calculated the ultimate creep strains and Poisson’s ratios (i.e., ***ϵ***_axial_, ***ϵ***_lateral_, and ν_app_) at the end of each creep cycle by averaging data from the last 20 seconds. We then conducted a two-way ANOVA followed by a multiple comparison Tukey’s test on each of ***ϵ***_axial_, ***ϵ***_lateral_, and ν_app_ with the anatomical region (I, N, S, T) and the creep cycle number (Cycles 1-4) as independent variables. The statistical significance threshold was set at 5%. Data visualization was performed using Gramm (Morel, 2018).

### 2.5 Finite element model of the ONH

#### 2.5.1 Mesh generation

To model deformation of the cylindrical ONH samples, we used an axisymmetric circular mesh subtending a 1 degree angle (Figure 1G). Consistent with the experimental samples’ dimensions, the height and the radius of the samples were each set to 0.5 mm. 220 linear elements were used to tessellate the geometry in 3D (Figure 1G); this mesh density was based on a preliminary convergence study (data not shown). The mesh was generated and the governing equations were solved using the FEBio software suite (FEBio v3.1; febio.org (Maas et al., 2012)).

#### 2.5.2 Boundary conditions

The boundary conditions were set to exploit the symmetry of the sample (Figure 1G). The rigid plate’s contact with the sample was implemented by a surface-to-surface penalty-based algorithm active both in tension and compression that allows for lateral deformation of the contact surface (Zimmerman and Ateshian, 2018). Because one of the contact bodies was a rigid plate, we assigned the contact surface from the deformable body (ONH) to be the primary contact surface and the plate’s contact surface as the secondary. The sample’s lateral surface was prescribed as a zero-pressure boundary surface to allow for fluid exchange during mechanical loading.

#### 2.5.3 Constitutive relations

To model the ONH’s mechanical behavior, we used a biphasic formulation with a viscoelastic matrix, similar to several existing studies considering other soft tissues (Ateshian, 2009; Ehlers et al., 2009; Huang et al., 2001). Here, we used a straightforward formulation of biphasic poroelasticity theory, using Darcy’s law to describe the solid-fluid interactions with an isotropic hydraulic permeability *k*, and a viscoelastic matrix to model the solid portion of the ECM using the reactive inelasticity (RIE) framework (Ateshian, 2015; Safa et al., 2019). RIE is a continuum mechanics framework based on the kinetics of molecular bonds to model the inelastic mechanical behaviors of tissue according to thermodynamics of observable state variables (Safa et al., 2019). In RIE’s viscoelastic variant (reactive viscoelasticity), there are two types of bonds: permanent bonds (also called strong bonds) and formative bonds (also called weak bonds) (Ateshian, 2015; Safa et al., 2019). Consequently, the Helmholtz free energy of the solid material (*Ψ*) can be written as

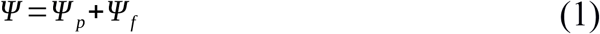

where *Ψ_p_* and *Ψ_f_* are the Helmholtz free energies of the permanent and formative bonds, respectively, and the subscripts ‘*p*’ and ‘*f*’ stand for permanent and formative, respectively. This notation will be used throughout the text.

Permanent bonds do not break-reform in response to external mechanical loading and thus confer hyperelastic mechanical behavior. In our model, *Ψ_p_* was formulated using a compressible neo-Hookean relation:

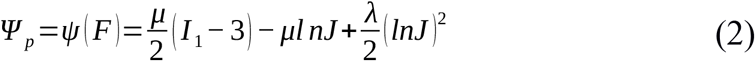

where *F* is the deformation gradient tensor, *I*_1_ is the first invariant of the right Cauchy-Green strain tensor (i.e., *I*_1_=*trace*(*C*)=*trace*(*F^T^ F*)), and *J* = *det*(*F*). Further, *μ* is the shear modulus (*μ*=*E/*[2(1+*ν*)]) and *λ* is the bulk modulus (*λ*=*Ev*/[(1+*ν*)(1 – 2*ν*)]) of the matrix, where *E* is the matrix Young’s modulus and *ν* is the matrix Poisson’s ratio.

Formative bonds, on the contrary, break-reform in response to external mechanical loading initiating a new generation of bonds (indicated by superscript α). The kinetics of these molecular bonds controls the amount of energy stored in them, resulting in viscoelastic behavior with a stress-free equilibrium condition. For this purpose *Ψ_f_* took the form

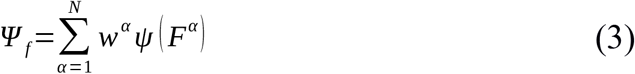

where *ψ* is the intrinsic hyperelasticity relation for formative bonds, which is taken to be the same as for permanent bonds (Eq. 2), *F^α^* is the relative deformation gradient tensor, and *w^α^* is the number fraction of a generation of bonds that was initiated in response to an increment of loading at time *t*=*t^α^*. *w^α^* is a scalar (0<*w^α^*<1), which obeys the conservation law (assuming no damage):

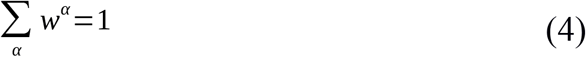

The kinetics of *w^α^* controls the transient part of the mechanical response; here we assumed first-order kinetics:

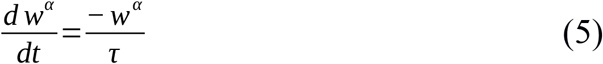

where *τ* is the time-constant of bond breaking-reforming. This form of the assumed kinetics results in an exponential stress-relaxation behavior (Ateshian, 2015; Safa et al., 2019).

In summary, there were a total of four model parameters: matrix Young’s modulus (*E* [kPa]), matrix Poisson’s ratio (*ν*), the time-constant of bond breaking-reforming (*τ* [sec]), and hydraulic permeability (*k* [mm^4^/(N.sec)]).

### 2.6 Parameter identification

To identify the model parameter values by fitting of experimental data, we used a multi-start optimization method (Safa et al., 2021, 2020). We based the optimization on a constrained nonlinear least-squares algorithm (interior-point, *fmincon*, Matlab) and used Latin hypercube sampling with a grid size of 100 to sample the initial guesses. The bounds of the model parameter values were {*E*, *ν*, *τ*, *k*}_min_= {0.1 kPa, 0, 10 sec, 10^-4^ mm^4^/(N.sec)} and {*E*, *ν*, *τ*, *k*}_max_= {100 kPa, 0.49, 1000 sec, 1 mm^4^/(N.sec)}.

The cost function was designed to fit the axial and lateral strains simultaneously, as:

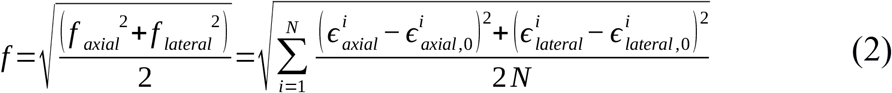

where *f_axial_* and *f_lateral_* are the root mean square of errors (RMSE) of the fits in the axial and lateral directions, respectively. ***ϵ***_axial,0_ and ***ϵ***_lateral,0_ are the experimental axial and lateral strains. The index *i* represents resampling intervals, which split the test duration into equal segments. We have previously shown that using both axial and lateral strains in data-fitting maximizes the parameter identifiability from uniaxial tests (Safa et al., 2021). Fits were preformed using a nonlinear constrained least-squares interior-point algorithm (*fmincon*, MATLAB), and the solutions were accepted if *f_axial_* and *f_lateral_* were less than 50% of the maximum experimental standard deviations of ***ϵ***_axial,0_ and ***ϵ***_lateral,0_, respectively (Safa et al., 2020). Additionally, only the solutions where the Hessian matrix was full rank with a positive determinant were accepted, indicating convexity of the function at the solution point (Hartmann et al., 2021). Due to the multi-start optimization’s stochastic nature, we conducted a confirmatory run, which resulted in the same results with minor, inconsequential differences that confirmed repeatability of the results.

### 2.7 Anatomical region heterogeneity in viscoelastic material properties

To assess the experimental variation in the viscoelastic properties of the ONH we conducted analyses of variance on each of the material parameters. Since the fit results were not Gaussian, we used a one-way non-parametric ANOVA test (Kruskal-Wallis test) for this purpose, followed with a conservative multiple comparison test using multiple Student’s t-tests with Bonferroni correction. A significance level of 5% was used for these analyses.

## 3 Results

### 3.1 Anatomical region heterogeneity in deformation responses

To assess possible effects of tissue heterogeneity, we analyzed the effect of anatomical region on the mechanical response of the ONH during compression testing, considering compression cycle number and anatomical region as independent variables (Figure 2). The anatomical region did not show an effect on axial strain (***ϵ***_axial_; p=0.7978); however, region was a significant factor for lateral strain (***ϵ***_lateral_; p=0.0091) and apparent Poisson ratio (ν_app_; p<0.0001). Specifically, for ***ϵ***_lateral_ the temporal region had a larger value compared to the nasal region (p<0.05), and for ν_app_ the temporal region had a larger value compared to both nasal and superior regions, but the inferior region only was larger relative to the nasal region (p<0.05; Figure 2D-F). As expected, cycle number had a strong effect on axial (p=0.0002) and lateral (p=0.0046) strains. However, ν_app_ was not affected by the cycle number (p=0.8224), i.e. ν_app_ was not dependent on the compression level. The p-values for the interaction between anatomical region and cycle number were almost unity for all the analyses, indicating that the anatomical heterogeneity was not affected by cycle number.

**Figure 2:**
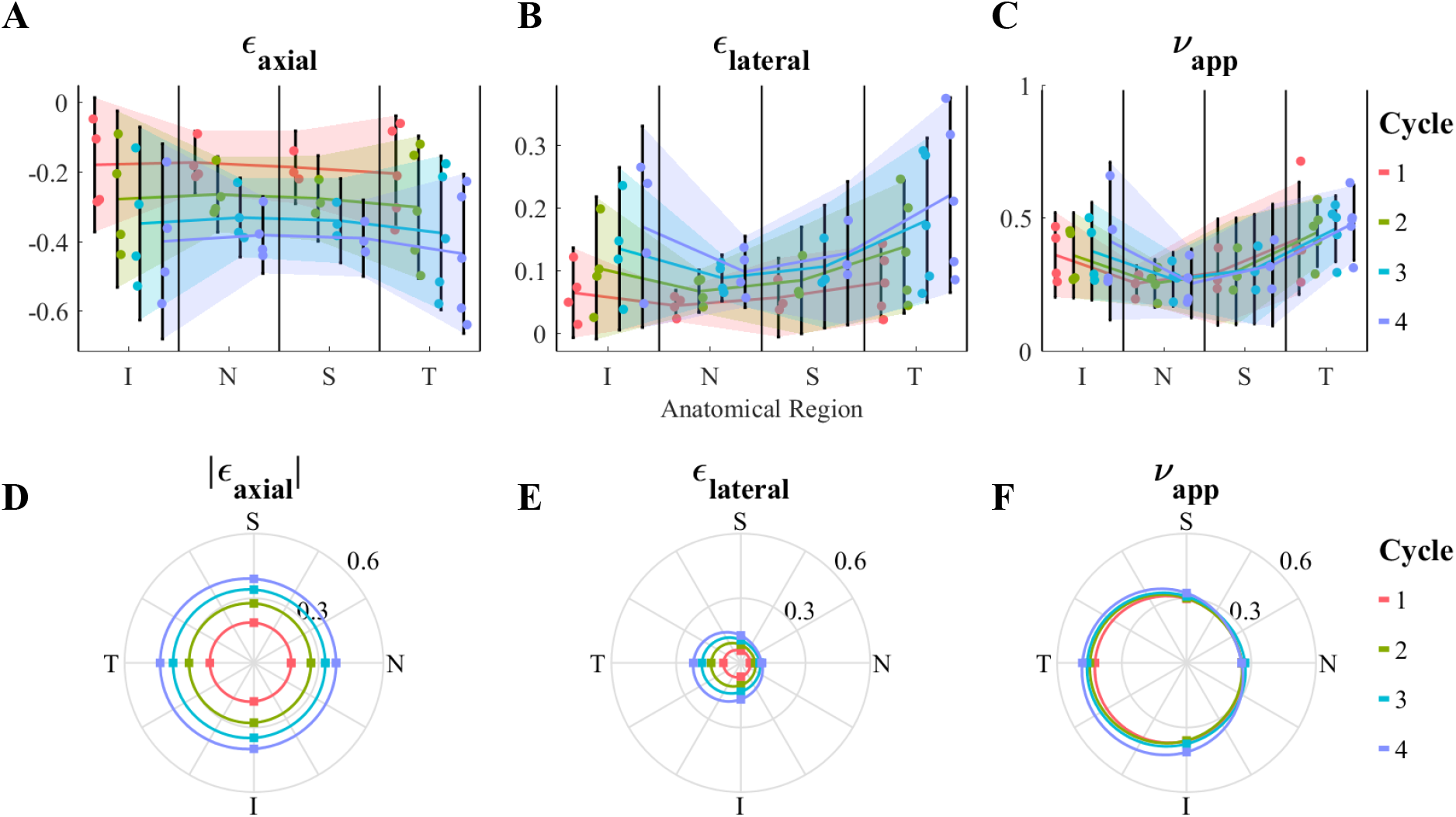
Regional dependence (analysis of variance) of (A and D) axial strain [*ϵ*_axial_], (B and E) lateral strain [*ϵ*_lateral_], and (C and F) apparent Poisson’s ratio [ν_app_]. For each parameter, the first row (A-C) shows the individual data points, their means, and 95% confidence intervals (shaded region) for each anatomical region and cycle number, and the second row (D-F) illustrates the mean values (dots) in a polar plot matching the anatomical orientations, where 0, 90, 180 and 270 degrees correspond to nasal [N], superior [S], temporal [T], and inferior [I], respectively. Although there was no significant effect of anatomical region on *ϵ*_axial_, both *ϵ*_lateral_ and ν_app_ were affected by the anatomical region, with the temporal and inferior quadrants having larger values compared to the nasal and superior quadrants. The cycle number significantly impacted *ϵ*_axial_ and *ϵ*_lateral_; however, it did not affect ν_app_. There was no interaction between the anatomical region and cycle number, i.e. heterogeneity was not affected by loading level.

### 3.2 Viscoelastic mechanical properties

The model fit the experimental data very well and captured both the axial and lateral strain behaviors for each of the anatomical regions with c. 75% of the fits passing the fit criteria (Figure 3). The best-fit parameter values for the ONH were *E*=1.895 [1.056,2 .391] kPa (median [min., max.]), *ν*=0.142 [0.060,0 .312], *τ*=213 [89,921] sec, and *k*=3.854 × 10^-1^ [3.457 × 10^-2^,9.994 × 10^-1^] mm^4^/(N sec) (Figure 4).

**Figure 3:**
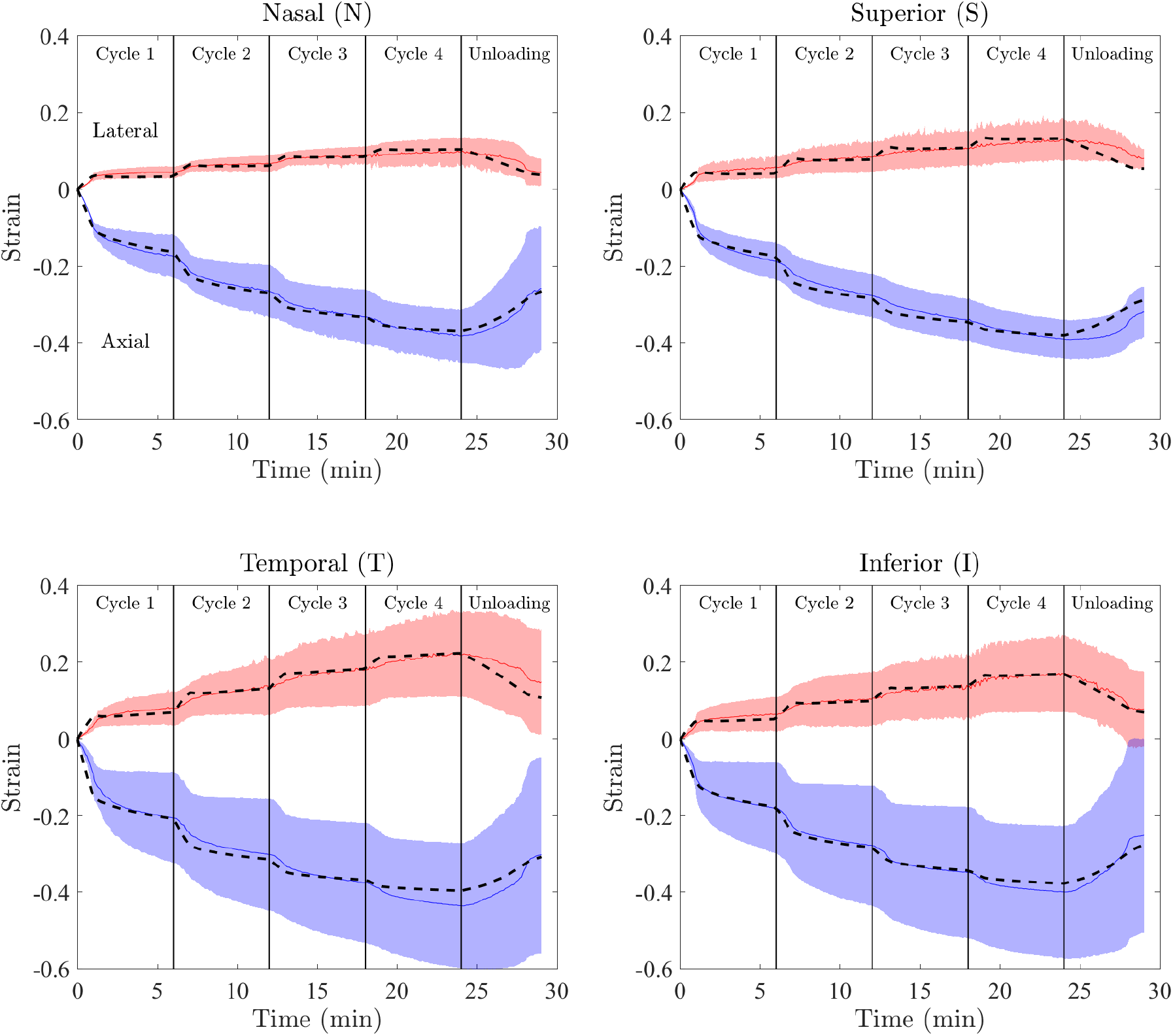
Results of data-fitting for each of the anatomical quadrants. The median and 95% confidence interval of the experimental data are shown with solid lines and shading, respectively. Positive strains correspond to the lateral direction, while negative values are axial strains. For each region, the model response generated using the median of accepted fits is shown by the dashed lines. Note the good agreement between experimental (solid lines) and fitted (dashed lines) strains.

**Figure 4:**
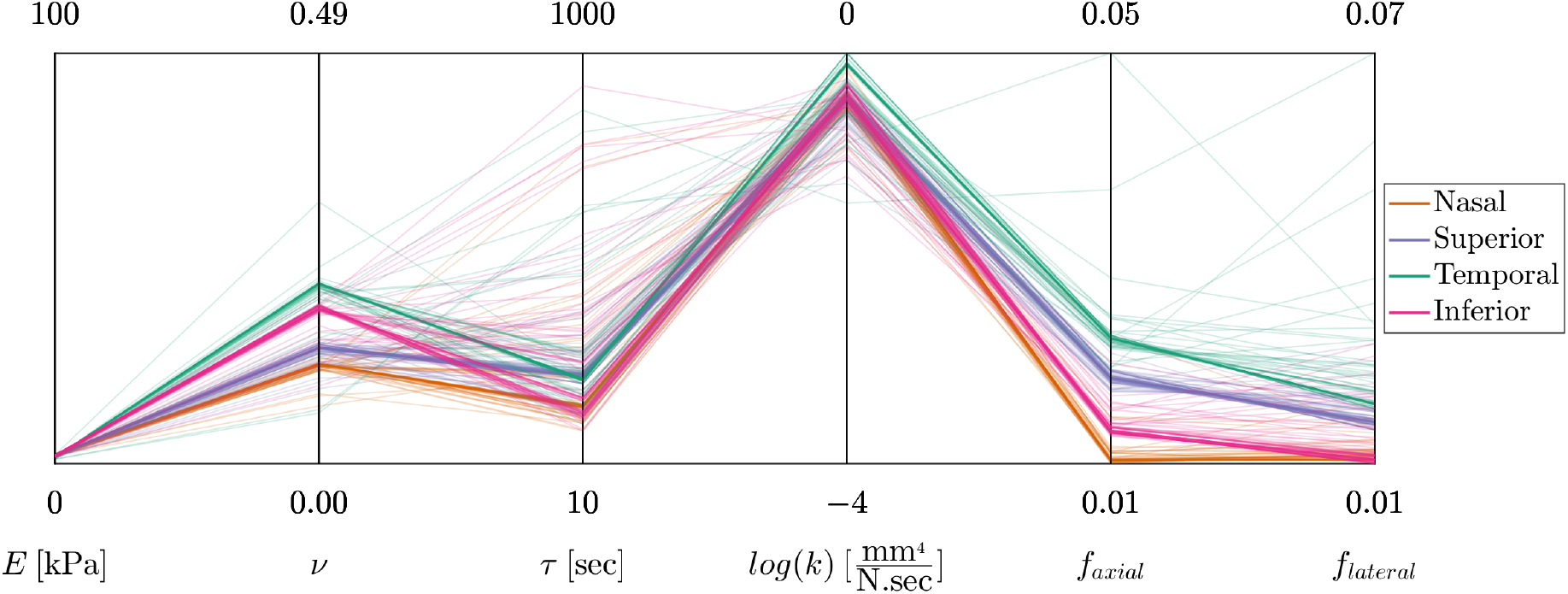
The data-fitting results shown in parallel coordinates format. The four coordinates on the left are the model parameters (*E*, *ν*, *τ*, *k*). For each model parameter, the graphed range of the coordinate corresponds to the prescribed range of model parameter values. The two additional coordinates are the RMSE of the fits in the axial and lateral direction (i.e., *f_axial_* and *f_lateral_*).

To assess the regional heterogeneity in the viscoelastic material properties, we plotted the fitted material parameter values and conducted analysis of variance (Figure 5). For each parameter, the median values had a nearly symmetrical distribution about the nasal-temporal axis, with either the nasal or temporal sides having the maximum/minimum values (Figure 5). In more detail, the nasal region (N) had the highest value of the Young’s modulus (Figure 5 A&E) compared to the rest of the regions, while the temporal region (T) had the lowest value. (Here and below, when a difference is cited it indicates statistical significance with p<0.05.) Conversely, the nasal region had the lowest value of Poisson’s ratio (*ν*) while the temporal region had the highest value (Figure 5 B&F). The variability was high for the time-constant (*τ*) with the distributions of the values of the regions being consistently skewed towards the range of 150-250 sec, and with the temporal and superior regions having slightly higher values (Figure 5 C&G). The permeability (*k*) value was highest in the temporal region (Figure 5 D&H).

**Figure 5:**
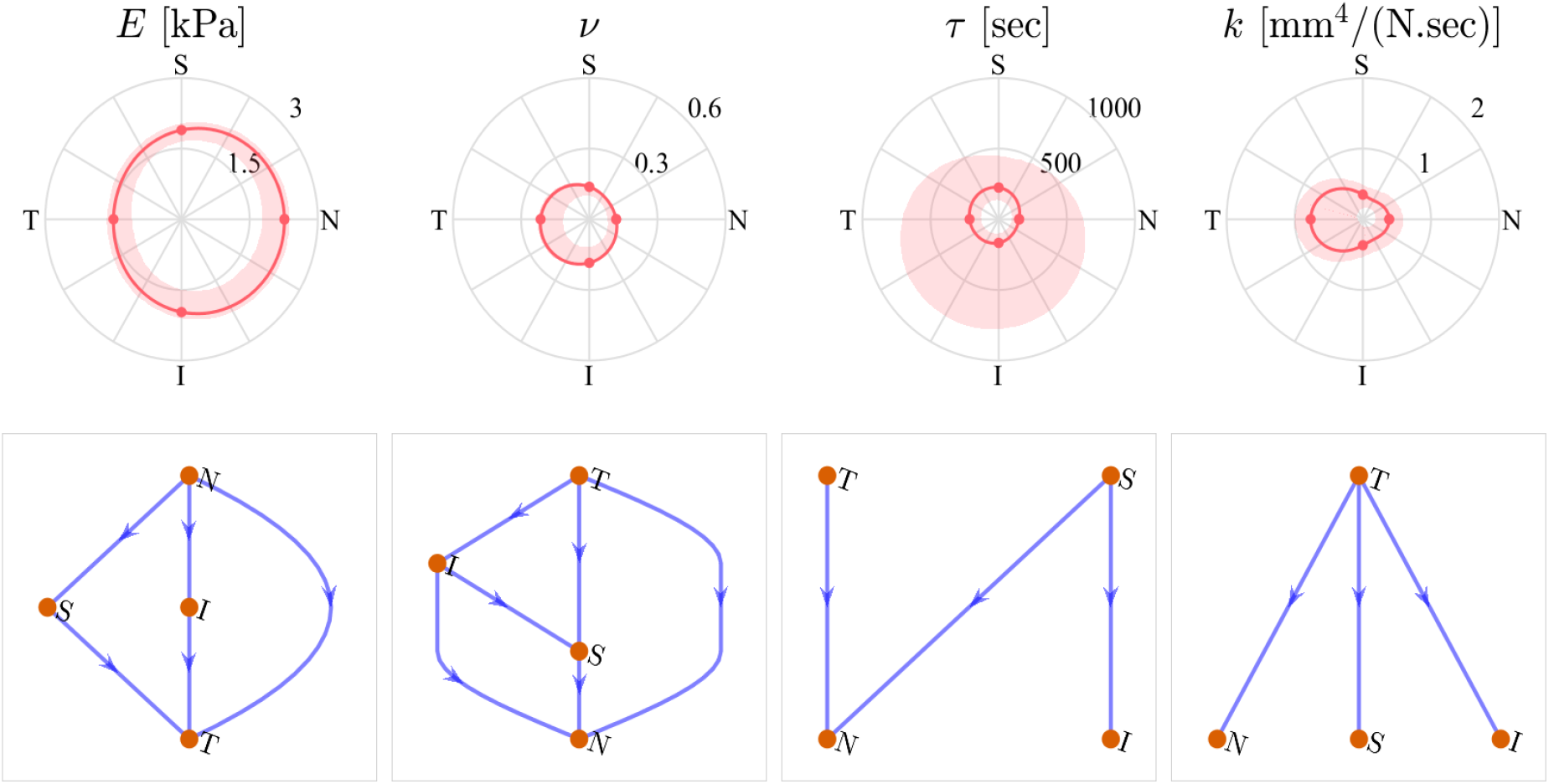
Anatomical distribution of the fit results. The first row shows the spatial distribution of each fitting parameter. The dots correspond to the median values, solid lines are the interpolated medians, and shaded areas are 95% confidence intervals for the acceptable fits (not the inter-sample variance). The second row shows the results of the multiple comparisons between the parameter values for each anatomical region, where every arrow connecting two regions (start and destination) indicate that the “start” region has a higher value compared to the “destination.”

## 4 Discussion

This study provides the first detailed evaluation of porcine optic nerve head (ONH) time-dependent biomechanical properties. Our results show that the ONH exhibits viscoelastic behavior, which can be quantitatively explained based on a combination of fluid-dependent and fluid-independent viscoelasticity. The former mechanism is a function of tissue porosity, while the latter is due to the biopolymeric structure of the ONH. Such a dual mechanism is common among other soft tissues exhibiting viscoelastic behavior (Connizzo and Grodzinsky, 2017; Huang et al., 2001).

Our fitted equilibrium Young’ modulus for the ONH was 1.896 [1.056, 2.391] kPa (median [min., max.]), which can be compared to previous quasistatic measurements on the ONH (3.05 kPa at 5% strain/min rate (Boazak et al., 2019)), and another study on the human lamina cribrosa (17.2 kPa (Brown et al., 2021)). Boazak and co-workers assumed that the ONH was incompressible (Boazak et al., 2019), which could explain the slightly larger modulus value they reported, since our experimental results showed that the ONH is compressible. More specifically, we found a fitted effective Poisson’s ratio *ν* of 0.142 [0.060, 0.312], and the apparent Poisson’s ratio was larger than *ν* (Figure 3B). This difference indicates the possibility of fluid exudation upon mechanical loading, and further illustrates the role of the fluid compression on the mechanical response of the tissue.

Compressibility is only one determinant of tissue’s fluid-dependent viscoelasticity, with the other principal factor being the hydraulic permeability, which controls the transient pressure build-up during tissue deformation. A small hydraulic permeability translates into larger pressurization. In this study, we report a porcine ONH hydraulic permeability of *k*=3.854 × 10^-1^ [3.457 × 10^-2^, 9.994 × 10^-1^] mm^4^/(N.sec), which is the first account of ONH hydraulic permeability we are aware of. Although not the same tissue as ONH, the porcine peripapillary sclera, which is anatomically adjacent, has a somewhat smaller permeability (*k*=0.86 × 10^-2^mm^4^/[N.sec]), and a stiffness of 10 kPa, which is larger than the stiffness we found for the ONH (Brown et al., 2021). The lower permeability and higher stiffness are consistent with the denser fibrous structure of the peripapillary sclera compared to the ONH (Boote et al., 2020; Jan et al., 2017).

The viscoelastic properties and the dynamic loading (e.g., rate of mechanical loading) are two important factors in tissue remodeling through controlling cellular mechanotransduction pathways (Suki et al., 2016). Our results indicate that the time constant of the solid matrix in the porcine ONH is in the range of 200 sec. Further, the median permeability of the ONH reported here corresponds to a biphasic characteristic time (*τ_bp_*=*a*^2^/(*H_a_k*), where *a* is the sample radius, *H_a_* is its aggregate modulus, and *k* is the hydraulic permeability (Armstrong et al., 1984)) of ~5.4 min for a sample with diameter of 1 mm (as tested in this study) and ~87 min for a sample with diameter of 4 mm, which is close to a full-size ONH (Figure 1D). This indicates that the ONH’s mechanical response is sensitive to dynamic loading with fluctuation times on the order of minutes to hours. This is an important observation because it indicates that ONH’s viscoelastic response is within the range of IOP dynamic fluctuations (from seconds [blinking] (Abelson, 2011) to hours [diurnal/nocturnal] (Kida et al., 2008, 2006)). These fluctuations may contribute to ONH remodeling by ONH cells, such as astrocytes and lamina cribrosa cells (Hernandez, 2000; Kirwan et al., 2005). Based on the time constants of the ONH it is likely that slower fluctuations (e.g., diurnal/nocturnal) are more relevant to remodeling in glaucoma than fast fluctuations, such as due to blinks, saccades and the ocular pulse; however, the IOP fluctuations occur at rates as high as 10,000 cycles per waking hour, which can also impact the effect of the slow and fast fluctuations (Turner et al., 2019). Further studies, such as 3D cell culturing and controlled mechanical loading of ONH cells is required to confirm this hypothesis.

Another interesting observation was the ONH’s heterogeneity, which can have important implications for understanding glaucoma pathophysiology. Our observations indicated that the nasal region had a higher stiffness and lower permeability (Figure 5), which was consistent with the observed axial strain responses, where the nasal region samples deformed less (Figure 2). Further, the nasal-temporal axis was an approximate axis of material property symmetry for the porcine ONH. Interestingly, in human ONH, a similar symmetry is observed where the region along the nasal-temporal axis has a denser collagenous structure compared to the rest of the ONH, strongly hinting at a similar trend in human ONH mechanical properties (Quigley and Addicks, 1981), which can be correlated with the patterns of the progression of glaucoma (Bartz-Schmidt et al., 1999). We note that although the symmetry in heterogeneity is consistent between porcine and human samples, it is unclear whether the relative differences in mechanical properties between the regions, especially between the nasal and temporal regions, are consistent between human and pigs. An interesting next step would be to use the framework established in this study to assess the human ONH’s mechanical properties. Such data could help further evaluate the fidelity of current animal models of ONH biomechanics, specifically by providing comparative information in addition to what is already known about posterior pole fibrous architecture (Gogola et al., 2018).

The loads imposed at each cycle of the mechanical test were motivated by physiological pressures, corresponding to approximately 5, 10, 15, and 20 mmHg acting on a circular surface area with diameter of 1mm. However, the resulting axial deformations were somewhat higher than those measured *in situ* in the porcine eye (Coudrillier et al., 2016). This is likely due to lack of peripapillary sclera tissue in our tests and the unconfined deformation boundary condition (Figure 1F). Although the strains and the unconfined compression boundary conditions do not exactly replicate the *in vivo* situation, they were necessary simplifications to enable direct experimental evaluation of the viscoelastic mechanical properties of the ONH at each anatomical quadrant of the ONH.

Our study was subject to some limitations. For example, we modeled the ONH as an isotropic material with deformation-independent permeability. This assumption is justifiable based on the lack of dependence of apparent Poisson’s ratio on compression level (Figure 2C); however, a strain-dependent permeability could improve the physiological relevance of the mathematical models. Further, we used large samples from each quadrant of the ONH (Figure 1D&E); this approach provides an average from each region, yet falls short of the spatial resolution required to appreciate the complex and fine laminar structure of the ONH. Atomic force microscopy has previously been used to assess the elastic properties of the ONH and lamina cribrosa [15], and a potential future approach to assess the viscoelastic mechanical response and its physical mechanisms in the ONH is to use high-bandwidth AFM-based rheology to provide higher spatial resolution (Connizzo and Grodzinsky, 2017; Oftadeh et al., 2018).

In conclusion, we evaluated ONH mechanical behavior using *in vitro* micromechanical testing and continuum mechanics modeling. We showed that the ONH’s viscoelastic mechanical response can be described by a combination of fluid-dependent and fluid-independent viscoelastic mechanisms. We also determined the viscoelastic mechanical properties of the porcine ONH for each of the four anatomical quadrants of the ONH. The material parameter values provided in this study can serve as benchmark values for future studies to replicate native tissue environments in *ex vivo* studies, such as using 3D cell cultures for studying mechanobiology and tissue engineering in glaucoma.

## 6 Acknowledgments

This study was supported by NIH R01 EY025286, and the Georgia Research Alliance. The authors thank T3 Labs (GCMI, Atlanta, GA) for the generous donation of the porcine eyes used in this study.

## References

Abelson, M.B., 2011. It’s Time to Think About the Blink. Review of Ophtalmology 58–61.

Armstrong, C.G., Lai, W.M., Mow, V.C., 1984. An Analysis of the Unconfined Compression of Articular Cartilage. Journal of Biomechanical Engineering 106, 165. https://doi.org/10.1115/1.3138475

Ateshian, G.A., 2015. Viscoelasticity using reactive constrained solid mixtures. Journal of Biomechanics 48, 941–947. https://doi.org/10.1016/j.jbiomech.2015.02.019

Ateshian, G.A., 2009. The role of interstitial fluid pressurization in articular cartilage lubrication. Journal of biomechanics 42, 1163–1176. https://doi.org/10.1016/j.jbiomech.2009.04.040

Bartz-Schmidt, K.U., Thumann, G., Jonescu-Cuypers, C.P., Krieglstein, G.K., 1999. Quantitative Morphologic and Functional Evaluation of the Optic Nerve Head in Chronic Open-Angle Glaucoma. Survey of Ophthalmology 44, S41–S53. https://doi.org/10.1016/S0039-6257(99)00076-4

Boazak, E.M., D’Humières, J., Read, A.T., Ethier, C.R., 2019. Compressive mechanical properties of rat and pig optic nerve head. Journal of Biomechanics 93, 204–208. https://doi.org/10.1016/j.jbiomech.2019.06.014

Boote, C., Sigal, I.A., Grytz, R., Hua, Y., Nguyen, T.D., Girard, M.J.A., 2020. Scleral structure and biomechanics. Progress in Retinal and Eye Research 74, 100773. https://doi.org/10.1016/j.preteyeres.2019.100773

Braunsmann, C., Hammer, C.M., Rheinlaender, J., Kruse, F.E., Schäffer, T.E., Schlötzer-Schrehardt, U., 2012. Evaluation of Lamina Cribrosa and Peripapillary Sclera Stiffness in Pseudoexfoliation and Normal Eyes by Atomic Force Microscopy. Invest. Ophthalmol. Vis. Sci. 53, 2960–2967. https://doi.org/10.1167/iovs.11-8409

Brown, D.M., Pardue, M.T., Ethier, C.R., 2021. A biphasic approach for characterizing tensile, compressive and hydraulic properties of the sclera. Journal of The Royal Society Interface 18, 20200634. https://doi.org/10.1098/rsif.2020.0634

Connizzo, B.K., Grodzinsky, A.J., 2017. Tendon exhibits complex poroelastic behavior at the nanoscale as revealed by high-frequency AFM-based rheology. Journal of Biomechanics 54, 11–18. https://doi.org/10.1016/j.jbiomech.2017.01.029

Ehlers, W., Karajan, N., Markert, B., 2009. An extended biphasic model for charged hydrated tissues with application to the intervertebral disc. Biomech Model Mechanobiol 8, 233–251. https://doi.org/10.1007/s10237-008-0129-y

Gogola, A., Jan, N.J., Lathrop, K.L., Sigal, I.A., 2018. Radial and circumferential collagen fibers are a feature of the peripapillary sclera of human, monkey, pig, cow, goat, and sheep. Investigative Ophthalmology and Visual Science 59, 4763–4774. https://doi.org/10.1167/iovs.18-25025

Hernandez, M.R., 2000. The optic nerve head in glaucoma: role of astrocytes in tissue remodeling. Progress in Retinal and Eye Research 19, 297–321. https://doi.org/10.1016/S1350-9462(99)00017-8

Huang, C.Y., Mow, V.C., Ateshian, G.A., 2001. The role of flow-independent viscoelasticity in the biphasic tensile and compressive responses of articular cartilage. Journal of biomechanical engineering 123, 410–417. https://doi.org/10.1115/1.1392316

Jan, N.-J., Lathrop, K., Sigal, I.A., 2017. Collagen Architecture of the Posterior Pole: High-Resolution Wide Field of View Visualization and Analysis Using Polarized Light Microscopy. Invest. Ophthalmol. Vis. Sci. 58, 735–744. https://doi.org/10.1167/iovs.16-20772

Kida, T., Liu, J.H.K., Weinreb, R.N., 2008. Effects of Aging on Corneal Biomechanical Properties and Their Impact on 24-hour Measurement of Intraocular Pressure. American Journal of Ophthalmology 146, 567–572. https://doi.org/10.1016/j.ajo.2008.05.026

Kida, T., Liu, J.H.K., Weinreb, R.N., 2006. Effect of 24-hour corneal biomechanical changes on intraocular pressure measurement. Investigative Ophthalmology and Visual Science 47, 4422–4426. https://doi.org/10.1167/iovs.06-0507

Kirwan, R.P., Fenerty, C.H., Crean, J., Wordinger, R.J., Clark, A.F., O’Brien, C.J., 2005. Influence of cyclical mechanical strain on extracellular matrix gene expression in human lamina cribrosa cells in vitro. Molecular vision 11, 798–810. https://doi.org/10.1016/s0021-9290(06)84552-5

Maas, S.A., Ellis, B.J., Ateshian, G.A., Weiss, J.A., 2012. FEBio: Finite Elements for Biomechanics. Journal of Biomechanical Engineering 134, 011005–011005. https://doi.org/10.1115/1.4005694

Martino, F., Perestrelo, A.R., Vinarský, V., Pagliari, S., Forte, G., 2018. Cellular mechanotransduction: From tension to function, Frontiers in Physiology. Frontiers Media S.A. https://doi.org/10.3389/fphys.2018.00824

Morel, P., 2018. Gramm: grammar of graphics plotting in Matlab. Journal of Open Source Software 3, 568. https://doi.org/10.21105/joss.00568

Myers, K.M., Cone, F.E., Quigley, H.A., Gelman, S., Pease, M.E., Nguyen, T.D., 2010. The in vitro inflation response of mouse sclera. Experimental Eye Research 91, 866–875. https://doi.org/10.1016/j.exer.2010.09.009

National Eye Institute, 2012. Vision Research: Needs, Gaps, and Opportunities. National Eye Institute 1–63.

Oftadeh, R., Connizzo, B.K., Nia, H.T., Ortiz, C., Grodzinsky, A.J., 2018. Biological connective tissues exhibit viscoelastic and poroelastic behavior at different frequency regimes: Application to tendon and skin biophysics. Acta Biomaterialia 70, 249–259. https://doi.org/10.1016/j.actbio.2018.01.041

Quigley, H.A., Addicks, E.M., 1981. Regional differences in the structure of the lamina cribrosa and their relation to glaucomatous optic nerve damage. Arch Ophthalmol 99, 137–143. https://doi.org/10.1001/archopht.1981.03930010139020

Ruiz-Ederra, J., García, M., Hernández, M., Urcola, H., Hernández-Barbáchano, E., Araiz, J., Vecino, E., 2005. The pig eye as a novel model of glaucoma. Experimental Eye Research 81, 561–569. https://doi.org/10.1016/j.exer.2005.03.014

Safa, Babak.N., Santare, Michael.H., Ethier, C.R., Elliott, Dawn.M., 2021. Identifiability of Tissue Material Parameters from Uniaxial Tests using Multistart Optimization. Acta Biomaterialia. https://doi.org/10.1016/j.actbio.2021.01.006

Safa, B.N., Bloom, E.T., Lee, A.H., Santare, M.H., Elliott, D.M., 2020. Evaluation of Transverse Poroelastic Mechanics of Tendon using Osmotic Loading and Biphasic Mixture Finite Element Modeling. Journal of Biomechanics 109, 109892. https://doi.org/10.1016/j.jbiomech.2020.109892

Safa, B.N., Santare, M.H., Elliott, D.M., 2019. A Reactive Inelasticity Theoretical Framework for Modeling Viscoelasticity, Plastic Deformation, and Damage in Fibrous Soft Tissue. Journal of Biomechanical Engineering 141, 1–12. https://doi.org/10.1115/1.4041575

Sanchez, I., Martin, R., Ussa, F., Fernandez-Bueno, I., 2011. The parameters of the porcine eyeball. https://doi.org/10.1007/s00417-011-1617-9

Suki, B., Parameswaran, H., Imsirovic, J., Bartolák-Suki, E., 2016. Regulatory roles of fluctuation-driven mechanotransduction in cell function, Physiology. American Physiological Society. https://doi.org/10.1152/physiol.00051.2015

Weinreb, R.N., Aung, T., Medeiros, F.A., 2014. The pathophysiology and treatment of glaucoma: A review, JAMA - Journal of the American Medical Association. American Medical Association. https://doi.org/10.1001/jama.2014.3192

World Health Organization, 2019. World report on vision, World health Organisation.

Zimmerman, B.K., Ateshian, G.A., 2018. A Surface-to-Surface Finite Element Algorithm for Large Deformation Frictional Contact in febio. Journal of Biomechanical Engineering 140, 081013–081013. https://doi.org/10.1115/1.4040497

